# A comparison between scalp- and source-reconstructed EEG networks

**DOI:** 10.1101/121764

**Authors:** Margherita Lai, Matteo Demuru, Arjan Hillebrand, Matteo Fraschini

## Abstract

EEG can be used to characterise functional networks using a variety of connectivity (FC) metrics. Unlike EEG source reconstruction, scalp analysis does not allow to make inferences about interacting regions, yet this latter approach has not been abandoned. Although the two approaches use different assumptions, conclusions drawn regarding the topology of the underlying networks should, ideally, not depend on the approach. The aim of the present work was to find an answer to the following questions: does scalp analysis provide a correct estimate of the network topology? how big are the distortions when using various pipelines in different experimental conditions? EEG recordings were analysed with amplitude- and phase-based metrics, founding a strong correlation for the global connectivity between scalp- and source-level. In contrast, network topology was only weakly correlated. The strongest correlations were obtained for MST leaf fraction, but only for FC metrics that limit the effects of volume conduction/signal leakage. These findings suggest that these effects alter the estimated EEG network organization, limiting the interpretation of results of scalp analysis. Finally, this study also suggests that the use of metrics that address the problem of zero lag correlations may give more reliable estimates of the underlying network topology.

## Introduction

Modern network science is a fundamental tool for the understanding of both normal and abnormal brain organization^1–3^. In this context, alongside other imaging techniques such as magnetoencephalography (MEG) and functional MRI (fMRI), electroencephalography (EEG) has been widely used to study brain networks^1,4^ It is well accepted that scalp-level EEG analysis does not allow inferences in terms of underlying neuroanatomy^5^, thus suggesting the use of source reconstruction techniques^6^, yet the former approach has not been abandoned. The main problems with scalp-level analysis of functional networks are related to two considerations: i) the already mentioned problem that location of EEG channels do not relate trivially to the location of the underlying sources; and ii) spurious estimates of functional connectivity can occur between the channels due to the effects of field spread^7^ and volume conduction^6^, where more than one channel can pick up the activity of an underlying source, although these effects can still be present in source-space^8^, where it is often referred to as signal leakage. Although scalp- and source-level analyses use different assumptions, conclusions drawn regarding the (global) topology of the underlying networks should, ideally, not depend on the approach that is used.

The aim of the present paper was to compare network measures extracted from scalp and source EEG signals, as defined by the minimum spanning tree (MST), and using a variety of functional connectivity (FC) metrics. The MST represents a network approach that provides an unbiased reconstruction of the core of a network^9,10^ MST parameters are sensitive to alterations in network topology^9^ and several studies have already implemented this approach in order to study network alteration at the scalp-^11–19^ and source-level^20–23^. Under certain conditions, the MST forms the critical backbone of the original network^24–26^. Moreover, it addresses several methodological limitations (i.e. biased estimates of network topology due to differences in connection strength or link density)^27^, yet still captures changes in topology of the original network^9^.

In order to assess the effect of field spread and volume conduction/leakage we included FC metrics that are sensitive to these effects, namely the phase locking value (PLV)^28^ and the amplitude envelope correlation (AEC)^29^, and metrics that are relatively insensitive to these effects, namely the phase lag index (PLI)^30^ and AEC after leakage correction^29,31^. Subsequently, several MST parameters were used to characterize the global topology of the reconstructed functional networks. We used a freely available EEG dataset^32,33^ that includes one minute eyes-closed and eyes-open resting-state recordings from 109 subjects. Source-level time-series of neuronal activity were reconstructed for 68 ROIs^34^ by means of the weighted Minimum Norm Estimate (wMNE)^35–38^ and segmented into five non-overlapping epochs. MSTs were reconstructed for four FC metrics, both at the scalp- and source-level, and parameters that characterise global topology were subsequently compared between the two domains. Finally, with the aim to examine the effect of using one of the two approaches (analysis at the scalp- or source-level) for different experimental conditions, and differently from previous works^39,40^ which adopted a simulation framework, we also show the results for both eyes-closed and eyes-open resting-state data.

## Material and methods

### Dataset

A dataset created by the developers of the BCI2000 instrumentation system (www.bci2000.org), consisting of 64-channels EEG recordings from 109 subjects, was used in this study^32,33^. All the EEG recordings analysed during the current study are available in the PhysioNet repository, https://www.physionet.org/pn4/eegmmidb/. The dataset contains fourteen different experimental runs for each subject, comprising of two one-minute baseline runs (eyes-open and eyes-closed resting-state conditions). For the aim of the present work, we considered both eyes-closed and eyes-open resting-state runs. The data are provided in EDF+ format and contain 64 raw EEG signals as per the international 10-10 system, sampled at 160 Hz.

### EEG pre-processing

EEGLAB (version 13_6_5b)^41^ was used to re-reference to common average reference and filter (with *fir 1* filter type) the EEG signals (band-pass filter between 1 and 70 Hz and 60 Hz notch filtering). Successively, ADJUST (version 1.1.1)^42^, a fully automatic algorithm based on Independent Component Analysis (ICA), was used to detect and remove artifacts from the filtered signals.

### Source reconstruction

Source-reconstructed time-series were obtained by using Brainstorm software (version 3.4)^43^. First, a head model was created using a symmetric boundary element method in Open-MEEG (version 2.3.0.1)^44,45^ based on the anatomy derived from the ICBM152 brain^46^. Time-series of neuronal activity were reconstructed using whitened and depth-weighted linear L2 minimum norm estimate (wMNE)^35–38^, with an identity matrix as noise covariance. Sources were constrained to the cortex and source orientation was perpendicular to the cortical surface^47^ To limit the effect of differences in network size^27^ between scalp- (64 channels) and source-analysis, source-reconstructed time-series were projected onto 68 regions of interest (ROIs) as defined by the Desikan-Killiany atlas^34^, where time-series for voxels within a ROI were averaged (after flipping the sign of sources with opposite directions).

### Connectivity metrics

Functional connectivity metrics that are either sensitive or insensitive to the effects of field spread and volume conduction/signal leakage, based on either amplitude or phase information, were used. In particular, we used AEC, a measure of amplitude coupling that uses linear correlations of the envelopes of the band-pass filtered signals^29,31^ and AEC_corrected_, a version that uses a symmetric orthogonalisation procedure to remove zero-lag correlations (implemented in the time domain^31^). Furthermore, we used the PLV^28^, a measure that quantifies the consistency of phase differences (including zero-lag), and the PLI^30^, a measure that quantifies the asymmetry of the distribution of phase differences between time series and that ignores zero-lag phase differences. The connectivity metrics were calculated for all epochs of each subject, after having band-pass filtered the scalp- or source-reconstructed time-series in the alpha band (8-13 Hz) and segmenting the one-minute recordings in five non-overlapping epochs of 12 seconds^48^.

### Functional Network Topology

The EEG channels (scalp-level analysis) and the atlas-based ROIs (source-level analysis) were considered as network nodes, and the functional connections as weighted edges within the network. Then the MST, a sub-network that connects all nodes whilst minimizing the link weights without forming loops, was reconstructed. Since we are interested in the strongest connections, the connection weights were inverted (1 – FC measure) before constructing the MST. The topology of the MST was characterised using the following parameters^51^: the leaf fraction (number of nodes with degree of 1 divided by the total number of nodes), the diameter (largest distance between any two nodes), the tree hierarchy (balance between hub overload and network integration) and the kappa (broadness of degree distribution). The use of MST was introduced as a simple and unbiased method to represent the essential features of brain networks. In particular, the MST, which captures the backbone of the network, allows to compare this feature across different conditions (i.e., behavioural states and neurological diseases). If the original network can be interpreted as a kind of transport network, and if edge weights in the original graph possess strong fluctuations, also called the strong disorder limit, all transport in the original graph flows over the MST^25^ forming the critical backbone of the original graph^52^. Moreover, it has been shown that the MST characterization still keeps important information about the topology of the whole network, as derived using more conventional graph analysis^9^. The MST leaf fraction captures if the network has a central organization, the diameter reflects the efficiency of the global organization, the tree hierarchy captures the balance between network efficiency and overload of central nodes (hubs), and kappa reflects the resilience of the network against attacks. A more detailed description and interpretation of MST parameters may be found in^9,10^. All analyses were performed using Matlab R2016b (The MathWorks, Inc., Natick, Massachusetts, US) and the MIT Strategic Engineering Matlab Tools for Network Analysis^53^.

### Statistical analysis

Correlations between scalp- and source-derived measures were assessed using Spearman’s rank correlation coefficient. In order to test differences between correlations from the different connectivity approaches, the ones that are insensitive to zero-lag correlations in contrast to the ones that are sensitive, we used a percentile bootstrap approach for non-overlapping correlations^54^, using 500 repetitions, and using the code available at https://github.com/GRousselet/blog/tree/master/comp2dcorr and described at https://garstats.wordpress.com/2017/03/01/comp2dcorr/.

## Results

The associations between measures extracted from scalp- and source-reconstructed networks are represented as scatterplot in Figure 1. The Spearman correlations between scalp- and source-level global connectivity (obtained by averaging all the values in the connectivity matrix excluding the diagonal) were high for all connectivity metrics, whereas correlations were low to moderate for the MST parameters. For the MSTs, the highest correlations were observed for leaf fraction for MSTs based on AEC_corrected_ (rho = 0.346) and PLI (rho = 0.262).

**Figure 1.**
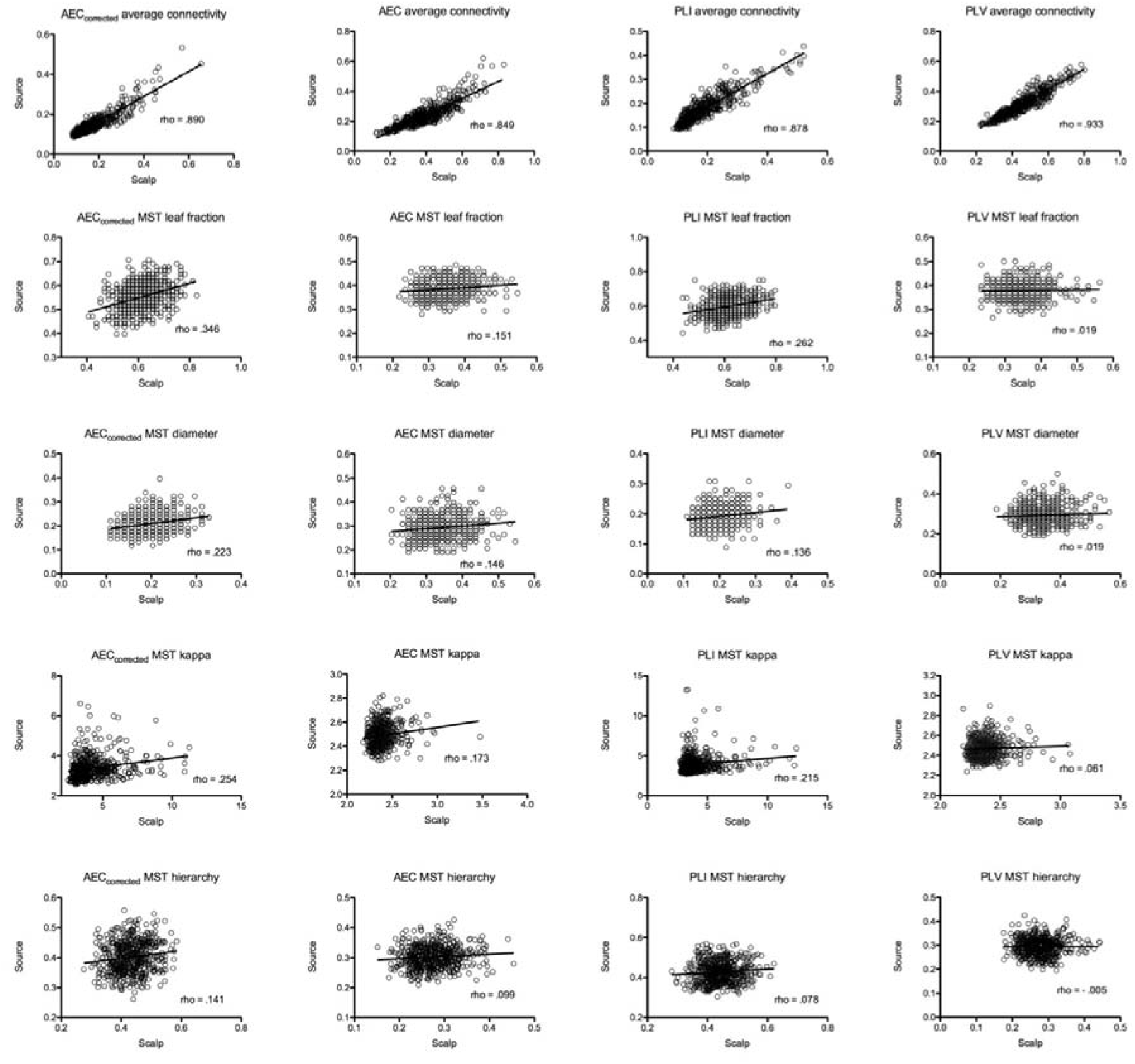
Scatterplots of scalp- and source-based measures of FC and network topology. The strength of the correlation is reported as rho value. Note that estimates of network topology correlated only weakly to moderately (maximum rho = 0.346) between scalp- and source-level, even though global FC correlated strongly.

For all the MST parameters, MSTs based on AEC_corrected_ and PLI showed higher correlations in comparison with AEC and PLV. In particular, PLV-based MST parameters showed the lowest correlations, with correlations strength approaching zero for all the evaluated network measures (maximum rho = 0.061, for MST kappa). Statistical differences, expressed using confidence intervals, between Spearman correlations derived from amplitude and phase based coupling approaches are summarized in Table1 and Table 2. For amplitude based FC metrics, the largest difference was observed for MST leaf fraction, whilst for the other MST parameters the differences were small. For phase-based FC metrics, the most marked differences were observed for MST leaf fraction and MST Kappa. For both the amplitude- and phase-based approaches, the MST Hierarchy showed only minimal differences. Figure 2 shows differences and bootstrap distributions for MST leaf fraction, for MSTs based on AEC_corrected_ versus AEC (left panel) and for MSTs based on PLI versus PLV (right panel).

**Table 1.**
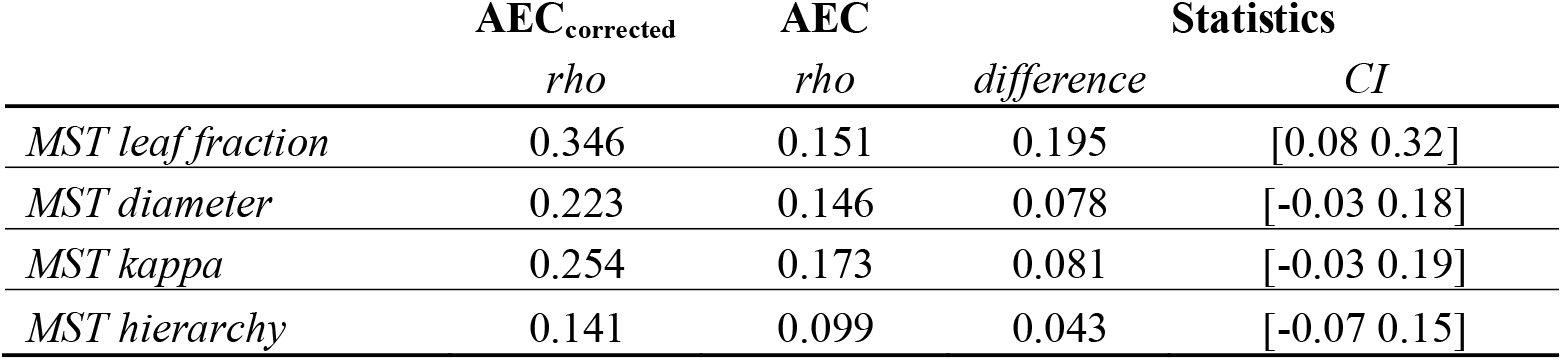
Comparison between scalp- and source-level correlations for amplitude based FC metrics. Mean difference and confidence intervals are reported.

**Table 2.** Comparison between scalp- and source-level correlations for phase based FC metrics. Mean difference and confidence intervals are reported.

|  | **PLI** | **PLV** | **Statistics** |  |
| --- | --- | --- | --- | --- |
|  | *rho* | *rho* | *difference* | *CI* |
| *MST leaf fraction* | 0.262 | 0.019 | 0.243 | [0.13 0.37] |
| *MST diameter* | 0.136 | 0.019 | 0.117 | [0.00 0.23] |
| *MST kappa* | 0.215 | 0.061 | 0.153 | [0.03 0.27] |
| *MST hierarchy* | 0.078 | -0.005 | 0.082 | [-0.04 0.19] |

**Figure 2.**
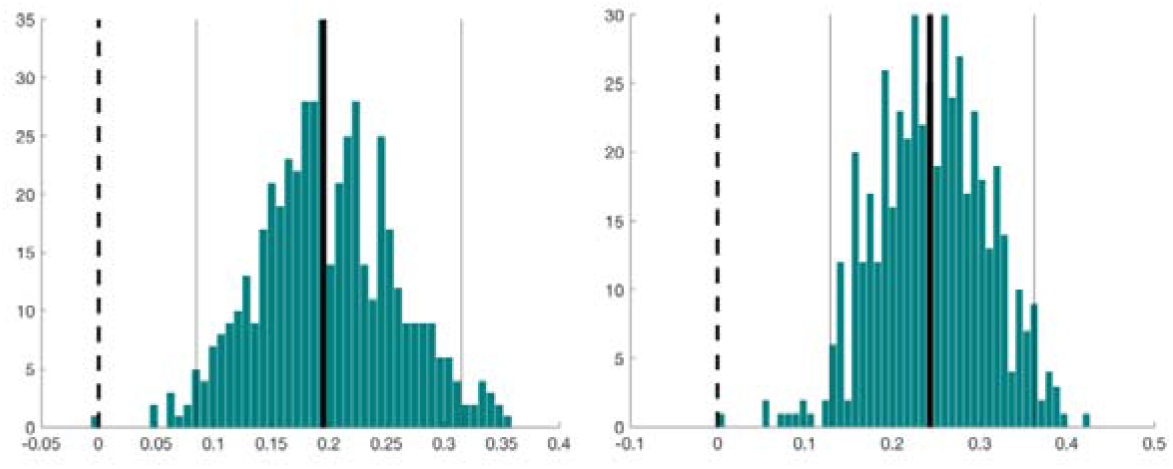
Differences and bootstrap distributions for MST leaf fraction, for AEC_corrected_ versus AEC (left panel) and for PLI versus PLV (right panel). The difference between coefficients is marked by a thick vertical black line. The 95% percentile bootstrap confidence interval is illustrated by the two thin vertical black lines.

As shown in Table 3, Spearman correlations tended to be higher (especially for PLI) when the same analysis was performed at subject-level. For this analysis, FC metrics from the five epochs were averaged for each single subject, and the correlation between scalp- and source-level estimates was computed.

**Table 3.** A Comparison between epoch-level and subject-level analysis. In the latter case, PLI-based correlations tended to be higher and in general outperformed AEC-based correlations.

|  | rho values |  |
| --- | --- | --- |
|  | Epoch level | Subject level |
| AEC global | 0.890 | 0.904 |
| PLI global | 0.878 | 0.918 |
| AEC MST leaf fraction | 0.346 | 0.325 |
| PLI MST leaf fraction | 0.262 | 0.367 |
| AEC MST diameter | 0.223 | 0.278 |
| PLI MST diameter | 0.136 | 0.253 |
| AEC MST kappa | 0.173 | 0.104 |
| PLI MST kappa | 0.215 | 0.181 |
| AEC MST hierarchy | 0.141 | 0.123 |
| PLI MST hierarchy | 0.078 | 0.151 |

Comparison of the two experimental conditions, namely eyes-closed and eyes-open resting-state, shows (Figure 3) that, for some connectivity metrics, adopting one approach over the other (scalp-versus source-level analysis) may show a different magnitude, and even an opposite direction, for the condition effect on network topology (here assessed by MST leaf fraction). It is interesting to note that, at least for those metrics that address the problem of field spread and volume conduction/signal leakage (AEC_corrected_ and PLI) the condition-related shift in network topology were in the same direction for the scalp- and sensor-level analysis, whereas for the other metrics (AEC and PLV) the condition-related shifts in network topology were in opposite directions, depending on whether the networks were reconstructed at the scalp- or source-level^55^.

**Figure 3.**
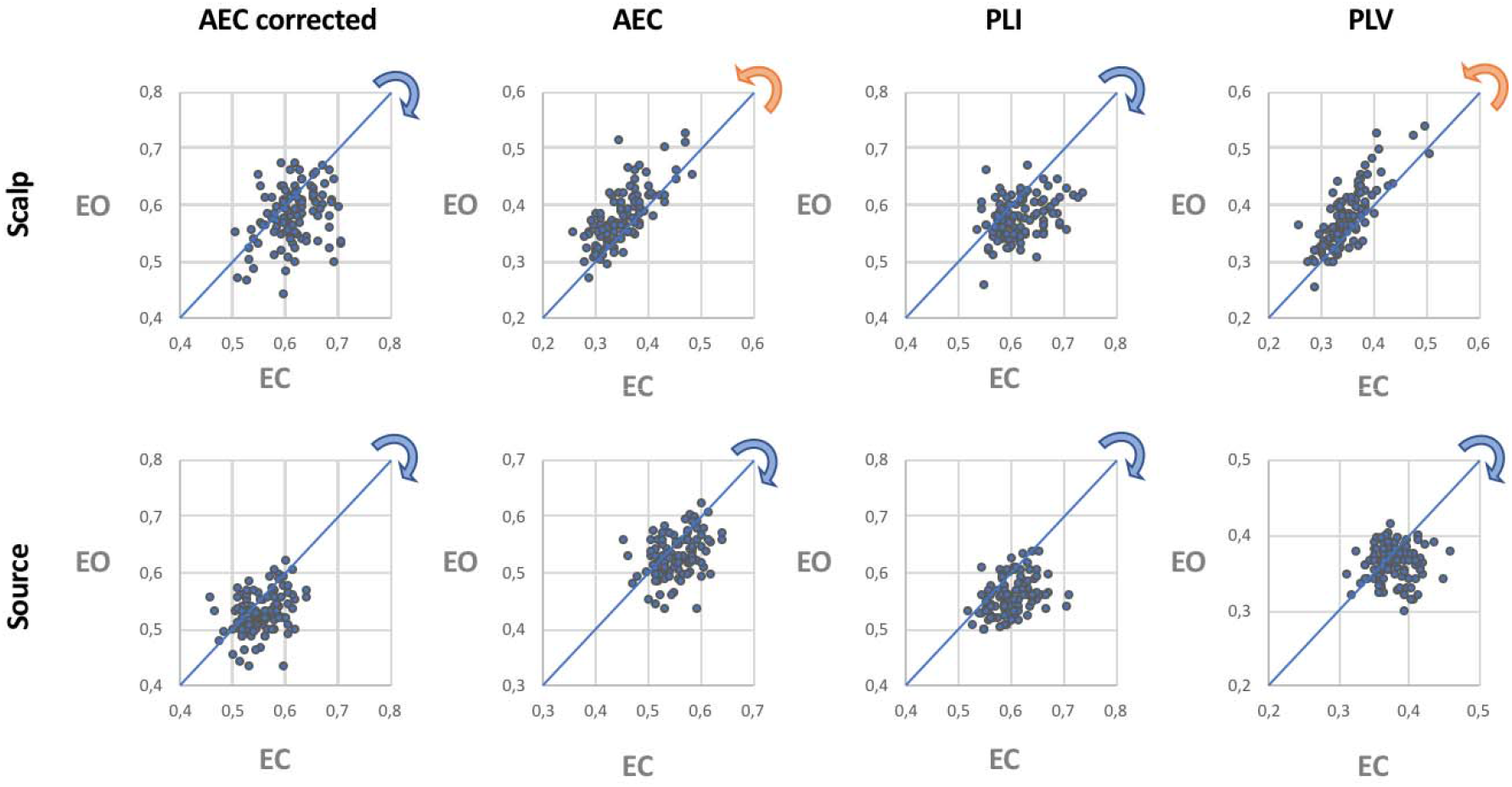
Scatterplot of leaf fraction for eyes-closed and eyes-open resting-state, based on scalp (top row) and source analysis (bottom row) and for each connectivity measure (different columns). The blue and orange arrows indicate the direction of the shift in network topology when comparing the eye-closed and eyes-open condition. Note that for the AEC and PLV the leaf fraction is higher in the eyes-open condition than in the eyes-closed condition for the scalp-level MSTs, whereas the opposite condition effect is found for the source-level MSTs.

## Discussion

Although network reconstructions at the scalp- and source-level rely on different assumptions, conclusions drawn regarding the (global) topology of the underlying networks should, ideally, not depend on the approach that is used. We found that global functional connectivity correlated strongly between the two domains, independent of the metric that was used. In contrast, global MST network descriptors extracted from scalp- and source-level EEG signals correlated (at best) moderately. In particular, in the case of connectivity metrics that do not limit spurious connections that are due to field spread and volume conduction/signal leakage (PLV and uncorrected version of AEC), the correlations were particularly weak. Although topological parameters for MSTs based on AEC_corrected_ and PLI showed only moderate correlations between scalp- and source-level, they were still higher than for MSTs based on PLV and uncorrected AEC for all of the estimated network measures. These differences (between scalp-/sensor-level correlations) were most evident for phase-based synchronization metrics, where PLI allowed to obtain higher correlations than PLV, for two of the MST descriptors (leaf fraction and kappa). These findings, which show minimal consistency between network analysis at scalp- and source-level, still advise against the use of FC metrics that do not correct for spurious correlations as they tend to amplify the differences between the two domains (scalp- and source-level). Conversely, the use of metrics that limit spurious connectivity (AEC_corrected_ and PLI) tends to reduce these differences. Higher correlations were observed when the analysis was performed at subject-level. Finally, the magnitude, and even direction, of condition-effects may change depending on the connectivity metric used (Figure 3), thus potentially leading to completely different conclusions. In this work, we used weighted minimum norm estimated to reconstruct the source activity.

It should be noted that different source reconstruction methods^56^, may provide different results in terms of global network topology and the correlation between scalp- and sensor-level estimates. In order to examine the potential effect of other inverse methods, we have replicated the analysis using the sLoreta approach^57^ which, as reported in the Supplementary Information, provided results, in terms of correlations between scalp- and source-level metrics, that were equivalent to those obtained with the wMNE. Since the accuracy of source localization may increase with the number of channels^39,58^, one would expect the correlations between networks reconstructed at the sensor- and source-level to be higher for MEG than for EEG. The reduced sensitivity of MEG volume conduction may further aid in this respect. It should also be noted that, differently from MEG, the choice of the reference electrode may influence EEG sensor-space connectivity estimates^59^. In this latter work^59^, the authors also report a detailed analysis about the effects of different approaches (such as anatomical templates, inverse methods and software implementations) on source-reconstructed EEG signals and their connectivity properties. Even though our work makes use of a set of different metrics to estimate functional connectivity and network topology, the reported results are consistent with those reported by Mahjoory et. al, who also suggest that the variability introduced by different methodological choices reflects uncertainty in the results, which should not be overlooked when interpreting scalp and source-level findings. It is also interesting to note that our findings seem to be in line with another recent work^60^ that investigated EEG coherence based on local field potentials measured with microelectrode arrays, as well as based on scalp-level EEG, and concluded that scalp-level coherence was not reliably related to coherence between brain areas measured intracranially.

The present work suffers from some limitations. First of all, there was no ground truth in our study, and we interpret our results under the assumption that network estimates at the source-level are a better approximation of the unknown true network organization than scalp-level estimates. This assumption is in line with a previous study^39^ that have highlighted that scalp-level network analyses may result in erroneous inferences about the underlying network topology. In particular, the model study by Antiqueira and colleagues suggests that scalp-based network structures, especially when under-sampled at surface sites, might not agree with the underlying threedimensional network. Another limitation refers to the inherently different mapping approaches between scalp- (channels) and source-level (ROIs) analysis, that strongly hinder the comparison between the two domains. Even though the number of EEG channels differed only slightly from the number of reconstructed ROIs (64 versus 68), it has been shown that differences in network size can affect estimates of network topology^27^. The correlations between the scalp- and source-level estimates of network topology presented here may therefore be lower than those that would have been obtained in case the networks had been of equal size. In order to estimate the possible effect of small differences in network size between sensor- and source-level we reported (see Supplementary Information) the results obtained using a subset of nodes for the source analysis. In particular, we omitted four nodes from original atlas (right and left parahippocampal and lingual) thus obtaining a network with 64 nodes. The results show that the impact of network size is negligible, since both the direction and the strength of associations (as measures by Spearman correlations, for MST leaf fraction based on AEC_corrected_) are comparable (difference = −0.01 [−0.03 0.01]). Another consideration is that consistency for local measures (e.g. nodal centrality) may be lower than reported for the global network measures. In order to investigate whether the observed correlations between scalp- and source-space based metrics were depended on the specific thresholding approach for the network reconstruction (i.e., the MST), we also replicated the analysis using a novel technique, namely efficiency cost optimization (ECO). This approach was recently introduced with the aim to filtering information in complex brain networks^61^. Our results show (see Supplementary Information) that also with ECO the functional connectivity metrics that are less sensitive to the effects of field spread and volume conduction/leakage (AEC_corrected_ and PLI) gave higher correlations between network metrics as obtained at scalp- and source-level, compared to AEC and PLV.

Although these results demonstrate that the specific thresholding approach does only have a small effect on the results for unweighted networks, it is unclear whether our results generalise to weighted and/or directed network analyses. Similarly, our results were obtained for specific connectivity metrics. We used metrics that are widely used, and that capture different mechanism of coupling between brain regions^49,50^, based on phase coupling or amplitude correlations. However, other metrics of functional connectivity may capture different properties of statistical interdependence, and may have a different sensitivity to, for example, common signal sources and non-stationarity. Different metrics may therefore have different biases when estimating interactions between time-series, which may also affect the correlation between source- and sensor-level results. Hence, although our findings indicate that results from sensor-level unweighted network analysis should be interpreted with caution, future studies should investigate whether this is also the case for weighted and/or directed networks that are derived from a larger range of functional connectivity metrics.

Moreover, we investigated the effect of frequency content (limited at alpha band in the main text), extending the analysis to other bands, namely theta (4 – 8 Hz) and beta (13 – 30 Hz) bands (see Supplementary Information). These results show that the reported trend still holds also for other frequency bands. Finally, the ICA-based artifact-rejection that we used may have distorted the phases^62^. However, since our study is focused on the comparison between analyses at the scalp- and source-level and is not intended to unveil the true underlying network topology, this limitation should only slightly impact on the reported results. Future studies should elucidate the effects of ICA artifact rejection on subsequent connectivity and network analyses.

## Conclusion

In conclusion, the present work confirms that, although functional connectivity can be estimated reliably, extreme caution should be used when interpreting results derived from scalp-level EEG network analysis, even when unbiased approaches such as MST analysis are used. However, assuming that the source-based network representation is a better approximation of the unknown true network organization, our findings also indicate that connectivity metrics that limit the emergence of spurious correlations (such as the corrected AEC and PLI) may allow for more reliable estimates of the underlying global network organization.

## Author Contributions

M. F., M. D. and A. H. conceived the study, M. L. and M. F. analysed the data, A. H., M. D., M. L. and M. F. wrote the draft of the paper. All authors reviewed the manuscript.

## Acknowledgements

None of the authors has any conflict of interest to disclose in relation to this work.

## Data availability

The datasets analysed during the current study are available in the PhysioNet repository, https://www.physionet.org/pn4/eegmmidb/.

## References

1. Stam, C. J. Modern network science of neurological disorders. Nat. Rev. Neurosci. 15, 683–95 (2014).

2. Bullmore, E. & Sporns, O. Complex brain networks: graph theoretical analysis of structural and functional systems. Nat. Rev. Neurosci. 10, 186–98 (2009).

3. Sporns, O., Chialvo, D. R., Kaiser, M. & Hilgetag, C. C. Organization, development and function of complex brain networks. Trends Cogn. Sci. 8, 418–425 (2004).

4. Stam, C. J. & van Straaten, E. C. W. The organization of physiological brain networks. Clin. Neurophysiol. 123, 1067–87 (2012).

5. Steen, F. Van de, Faes, L., Karahan, E. & Songsiri, J. Critical comments on EEG sensor space dynamical connectivity analysis. Brain Topogr. Epub ahead of print (2016).

6. Schoffelen, J.-M. & Gross, J. Source connectivity analysis with MEG and EEG. Hum. Brain Mapp. 30, 1857–1865 (2009).

7. Dominguez, L. G., Wennberg, R., Velazquez, J. L. P. & Erra, R. G. Enhanced measured synchronization of unsynchronized sources: inspecting the physiological significance of synchronization analysis of whole brain electrophysiological recordings. Int. J. Phys. Sci. 2, 305–317 (2007).

8. Brookes, M., Woolrich, M. & Price, D. An Introduction to MEG connectivity measurements. in Magnetoencephalography 321–358 (Springer Berlin Heidelberg, 2014).

9. Tewarie, P., van Dellen, E., Hillebrand, A. & Stam, C. J. The minimum spanning tree: An unbiased method for brain network analysis. Neuroimage 104, 177–188 (2015).

10. Stam, C. J. et al. The trees and the forest: Characterization of complex brain networks with minimum spanning trees. Int. J. Psychophysiol. 92, 129–138 (2014).

11. Vourkas, M. et al. Simple and difficult mathematics in children: A minimum spanning tree EEG network analysis. Neurosci. Lett. 576, 28–33 (2014).

12. Yu, M. et al. Different functional connectivity and network topology in behavioral variant of frontotemporal dementia and Alzheimer’s disease: an EEG study. Neurobiol. Aging 42, 150–162 (2016).

13. van Diessen, E., Otte, W. M., Stam, C. J., Braun, K. P. J. & Jansen, F. E. Electroencephalography based functional networks in newly diagnosed childhood epilepsies. Clin. Neurophysiol. 127, 2325–2332 (2016).

14. Fraga González, G. et al. Graph analysis of EEG resting state functional networks in dyslexic readers. Clin. Neurophysiol. 127, 3165–3175 (2016).

15. Fraschini, M. et al. The re-organization of functional brain networks in pharmaco-resistant epileptic patients who respond to VNS. Neurosci. Lett. 580, 153–7 (2014).

16. Fraschini, M. et al. EEG functional network topology is associated with disability in patients with amyotrophic lateral sclerosis. Sci. Rep. 6, 38653 (2016).

17. Demuru, M., Fara, F. & Fraschini, M. Brain network analysis of EEG functional connectivity during imagery hand movements. J. Integr. Neurosci. 12, 441–7 (2013).

18. Crobe, A., Demuru, M., Didaci, L., Marcialis, G. L. & Fraschini, M. Minimum spanning tree and k-core decomposition as measure of subject-specific EEG traits. Biomed. Phys. Eng. Express 2, 017001 (2016).

19. Fraschini, M., Hillebrand, A., Demuru, M., Didaci, L. & Marcialis, G. L. An EEG-Based Biometric System Using Eigenvector Centrality in Resting State Brain Networks. IEEE Signal Process. Lett. 22, 666–670 (2015).

20. Tewarie, P. et al. Functional brain network analysis using minimum spanning trees in Multiple Sclerosis: an MEG source-space study. Neuroimage 88, 308--318 (2014).

21. Dubbelink, K. T. E. O. et al. Disrupted brain network topology in Parkinson’s disease□: a longitudinal magnetoencephalography study. Brain 197–207 (2014). doi:10.1093/brain/awt316

22. van Dellen, E. et al. Epilepsy surgery outcome and functional network alterations in longitudinal MEG: a minimum spanning tree analysis. Neuroimage 86, 354–63 (2014).

23. Nissen, I. A. et al. Identifying the epileptogenic zone in interictal resting-state MEG source-space networks. Epilepsia 58, 137–148 (2017).

24. Wang, H., Hernandez, J. M. & Van Mieghem, P. Betweenness centrality in a weighted network. Phys. Rev. E 77, 046105 (2008).

25. Van Mieghem, P. & van Langen, S. Influence of the link weight structure on the shortest path. Phys. Rev. E 71, 056113 (2005).

26. Van Mieghem, P. & Magdalena, S. M. Phase transition in the link weight structure of networks. Phys. Rev. E 72, 056138 (2005).

27. van Wijk, B. C. M., Stam, C. J. & Daffertshofer, A. Comparing brain networks of different size and connectivity density using graph theory. PLoS One 5, e13701 (2010).

28. Lachaux, J. P., Rodriguez, E., Martinerie, J. & Varela, F. J. Measuring phase synchrony in brain signals. Hum. Brain Mapp. 8, 194–208 (1999).

29. Brookes, M. J. et al. Measuring functional connectivity using MEG: Methodology and comparison with fcMRI. Neuroimage 56, 1082–1104 (2011).

30. Stam, C. J., Nolte, G. & Daffertshofer, A. Phase lag index: assessment of functional connectivity from multi channel EEG and MEG with diminished bias from common sources. Hum. Brain Mapp. 28, 1178–93 (2007).

31. Hipp, J. F., Hawellek, D. J., Corbetta, M., Siegel, M. & Engel, A. K. Large-scale cortical correlation structure of spontaneous oscillatory activity. Nat. Neurosci. 15, 884–890 (2012).

32. Schalk, G., McFarland, D. J., Hinterberger, T., Birbaumer, N. & Wolpaw, J. R. BCI2000: a general-purpose brain-computer interface (BCI) system. IEEE Trans. Biomed. Eng. 51, 1034–43 (2004).

33. Goldberger, A. L. et al. PhysioBank, PhysioToolkit, and PhysioNet : Components of a New Research Resource for Complex Physiologic Signals. Circulation 101, e215–e220 (2000).

34. Desikan, R. S. et al. An automated labeling system for subdividing the human cerebral cortex on MRI scans into gyral based regions of interest. Neuroimage 31, 968–980 (2006).

35. Hämäläinen, M. S. & Ilmoniemi, R. J. Interpreting magnetic fields of the brain: minimum norm estimates. Med. Biol. Eng. Comput. 32, 35–42 (1994).

36. Hämäläinen, M. S. Interpreting measured magnetic fields in the brain: Estimates of current distributions. (1984).

37. Fuchs, M., Wagner, M., Köhler, T. & Wischmann, H. A. Linear and nonlinear current density reconstructions. J. Clin. Neurophysiol. 16, 267–95 (1999).

38. Lin, F.-H. et al. Assessing and improving the spatial accuracy in MEG source localization by depth-weighted minimum-norm estimates. Neuroimage 31, 160–171 (2006).

39. Antiqueira, L., Rodrigues, F. A., van Wijk, B. C. M., Costa, L. da F. & Daffertshofer, A. Estimating complex cortical networks via surface recordings—A critical note. Neuroimage 53, 439–449 (2010).

40. Anzolin, A. et al. Effect of head volume conduction on directed connectivity estimated between reconstructed EEG sources. bioRxiv 251223 (2018). doi:10.1101/251223

41. Delorme, A. & Makeig, S. EEGLAB: an open source toolbox for analysis of single-trial EEG dynamics including independent component analysis. J. Neurosci. Methods 134, 9–21 (2004).

42. Mognon, A., Jovicich, J., Bruzzone, L. & Buiatti, M. ADJUST: An automatic EEG artifact detector based on the joint use of spatial and temporal features. Psychophysiology 48, 229–240 (2011).

43. Tadel, F., Baillet, S., Mosher, J. C., Pantazis, D. & Leahy, R. M. Brainstorm: a user-friendly application for MEG/EEG analysis. Comput. Intell. Neurosci. 2011, 879716 (2011).

44. Kybic, J. et al. A common formalism for the Integral formulations of the forward EEG problem. IEEE Trans. Med. Imaging 24, 12–28 (2005).

45. Gramfort, A., Papadopoulo, T., Olivi, E. & Clerc, M. OpenMEEG: opensource software for quasistatic bioelectromagnetics. Biomed. Eng. Online 9, 45 (2010).

46. Mazziotta, J. et al. A four-dimensional probabilistic atlas of the human brain. J. Am. Med. Inform. Assoc. 8, 401–30 (2001).

47. Mosher, J. C., Leahy, R. M. & Lewis, P. S. EEG and MEG: forward solutions for inverse methods. IEEE Trans. Biomed. Eng. 46, 245–59 (1999).

48. Fraschini, M. et al. The effect of epoch length on estimated EEG functional connectivity and brain network organisation. J. Neural Eng. 13, 036015 (2016).

49. Engel, A. K., Gerloff, C., Hilgetag, C. C. & Nolte, G. Intrinsic Coupling Modes: Multiscale Interactions in Ongoing Brain Activity. Neuron 80, 867–886 (2013).

50. Guggisberg, A. G. et al. Two intrinsic coupling types for resting-state integration in the human brain. Brain Topogr. 28, 318–29 (2015).

51. Boersma, M. et al. Growing trees in child brains: graph theoretical analysis of electroencephalography-derived minimum spanning tree in 5- and 7-year-old children reflects brain maturation. Brain Connect. 3, 50–60 (2013).

52. Van Mieghem, P. & Magdalena, S. M. Phase transition in the link weight structure of networks. Phys. Rev. E - Stat. Nonlinear, Soft Matter Phys. 72, 1–7 (2005).

53. Bounova, G. & de Weck, O. Overview of metrics and their correlation patterns for multiple-metric topology analysis on heterogeneous graph ensembles. Phys. Rev. E 85, 016117 (2012).

54. Wilcox, R. R. Comparing dependent robust correlations. Br. J. Math. Stat. Psychol. 69, 215–224 (2016).

55. Rousselet, G. A., Foxe, J. J. & Bolam, J. P. A few simple steps to improve the description of group results in neuroscience. Eur. J. Neurosci. 44, 2647–2651 (2016).

56. Baillet, S., Mosher, J. C. & Leahy, R. M. Electromagnetic brain mapping. IEEE Signal Process. Mag. 18, 14–30 (2001).

57. Pascual-Marqui, R. D. Standardized low-resolution brain electromagnetic tomography (sLORETA): technical details. Methods Find. Exp. Clin. Pharmacol. 24 Suppl D, 5–12 (2002).

58. Lantz, G., Grave de Peralta, R., Spinelli, L., Seeck, M. & Michel, C.. Epileptic source localization with high density EEG: how many electrodes are needed? Clin. Neurophysiol. 114, 63–69 (2003).

59. Mahjoory, K. et al. Consistency of EEG source localization and connectivity estimates. Neuroimage 152, 590–601 (2017).

60. Snyder, A. C., Issar, D. & Smith, M. A. What does scalp electroencephalogram coherence tell us about long-range cortical networks? Eur. J. Neurosci. (2018). doi:10.1111/ejn.13840

61. De Vico Fallani, F., Latora, V. & Chavez, M. A Topological Criterion for Filtering Information in Complex Brain Networks. PLOS Comput. Biol. 13, e1005305 (2017).

62. Castellanos, N. P. & Makarov, V. A. Recovering EEG brain signals: Artifact suppression with wavelet enhanced independent component analysis. J. Neurosci. Methods 158, 300–312 (2006).

